# SPAGHETTI leverages massive H&E morphological models for phase contrast microscopy images with a generative deep learning approach

**DOI:** 10.1101/2025.08.27.672636

**Authors:** Zhi Fei Dong, Chris McIntosh, Gregory W. Schwartz

## Abstract

Phase contrast microscopy (PCM) is a powerful cell imaging method, one of the few technologies to delineate and track cell structure in live cells without staining. Despite PCM’s great potential and popularity in monitoring live-cell populations, there is a lack of algorithms to extract morphological information from these images due to the lack of large training datasets. To overcome this challenge and enable advanced, high throughput quantitative analysis of PCM images, we introduce SPAGHETTI: a lightweight image translator built on a modified cycle-consistent generative adversarial network. SPAGHETTI translates PCM images into those resembling hematoxylin and eosin (H&E) pathological images which, due to the pervasive and widespread use in clinical settings, are the basis for most large-scale deep learning models for quantitative analyses. We demonstrate that by first using SPAGHETTI to translate PCM images into H&E-like images, we could achieve significantly improved performance on cell segmentation through the use of tissue and H&E-specific cell segmentation models. We also show that by passing translated PCM images across several independent datasets into H&E feature extractor models, we improve the performance of cell-type annotation, experimental media classification, and cell viability prediction. Overall, SPAGHETTI enables many quantitative analyses of PCM that were previously impossible and acts as a valuable preprocessing step to help researchers gather novel information about cell states through the downstream quantitative analysis of morphological features.

## Introduction

Cell morphology is fundamental to understanding cell state and function. For example, cells respond to different stimuli and stress by changing their morphology to best adapt to the new environment and initiate proper cellular responses^1^. Cell morphology is also closely related to the cell’s pathological conditions, as irregular cell shapes and sizes are often associated with diseases such as cancer^2^. Therefore, the ability to observe and analyze cell morphology is crucial for discovering and understanding the cellular mechanisms underlying various pathological conditions, which can lead to the development of new diagnostic and therapeutic strategies^3^.

To characterize cell morphologies, researchers often rely on imaging techniques such as microscopy. Among the many microscopy techniques available, phase contrast microscopy (PCM) is an old yet widely used technique in modern in vitro cell biology research. Compared to other non-stained methods such as bright-field microscopy, PCM provides a more detailed and contrasted image of cells by converting the phase shift of the light passing through cells into brightness changes^4^. PCM’s ability to visualize the internal structures of cells without the need of staining also makes PCM a popular choice for quick live-cell imaging and temporal cell culture monitoring as long time monitoring of fluorescence stains are known to cause phototoxicity^5^. However, despite the popularity of PCM in cell state monitoring, these images are rarely used in analysis routines by biologists. Although PCM images can provide a plethora of information describing cells, few attempts have been made to extract this information quantitatively, forcing biologists to rely on assay measurements and readings from stain intensities. The lack of PCM usage not only adds additional steps and cost to experiments, but also limits the amount of information that can be extracted from images as assays and stain intensities can only measure a very limited number of properties at once.

Pathologists previously faced similar problems analyzing histopathology images. Hema-toxylin and eosin (H&E) staining is the most common staining method used in pathology to visualize cell morphology and provides a wealth of information characterizing tissue disease state. Until recently, the main method of H&E feature extraction was manual char-acterization by eye, and the analysis of H&E images was often qualitative and subjective^6^. In an effort to automate this process, recent advances in deep learning have lead to the development of high-performing foundational cell segmentors^7,8^ and feature extractors^9–11^ to segment cells and extract features from H&E images for downstream tasks such as diagnosis and survival prediction^12–14^, suggesting the important role of deep learning to extract a plethora of information from cellular morphologies to more quantitatively analyze H&E images. Despite these advances, cells from H&E assays result in static cells frozen in time, necessitating the need for techniques such as PCM. However, we cannot train the same foundational models for PCM images. Unlike H&E images that can be found in rich resources such as The Cancer Genome Atlas^15^, the lack of publicly available PCM datasets prevents the development of a high performing foundational morphological model which requires a large amount of data to train.

To overcome this challenge, we developed *SPAGHETTI (Ssim-restrained PhAse con-trast Gan to H&E TranslaTion of Images)*, a generative model that leverages massive H&E morphological models to improve tasks such as cell segmentation, classification, and regression for PCM images by translating them to H&E-like images (Figure 1a). Ar-chitectures such as cycle-consistent generative adversarial network (CycleGAN)^16^ can translate images from one domain to another without the need of paired images, applicable to various tasks in the field of medical image translation^17,18^. SPAGHETTI expands upon the original CycleGAN architecture that uses the cycle-consistent property in translation to both translate PCM images to H&E-like images and vice versa by introducing an additional Structure Similarity Index Measurement (SSIM) loss to eliminate hallucinations. Here, we demonstrate that by using SPAGHETTI as a preprocessing step, we can leverage founda-tional tissue and H&E segmentors and feature extractors for use on PCM images for cell segmentation and other quantitative downstream classification tasks. SPAGHETTI is open source and freely available at https://github.com/schwartzlab-methods/spaghetti.

**Figure 1:**
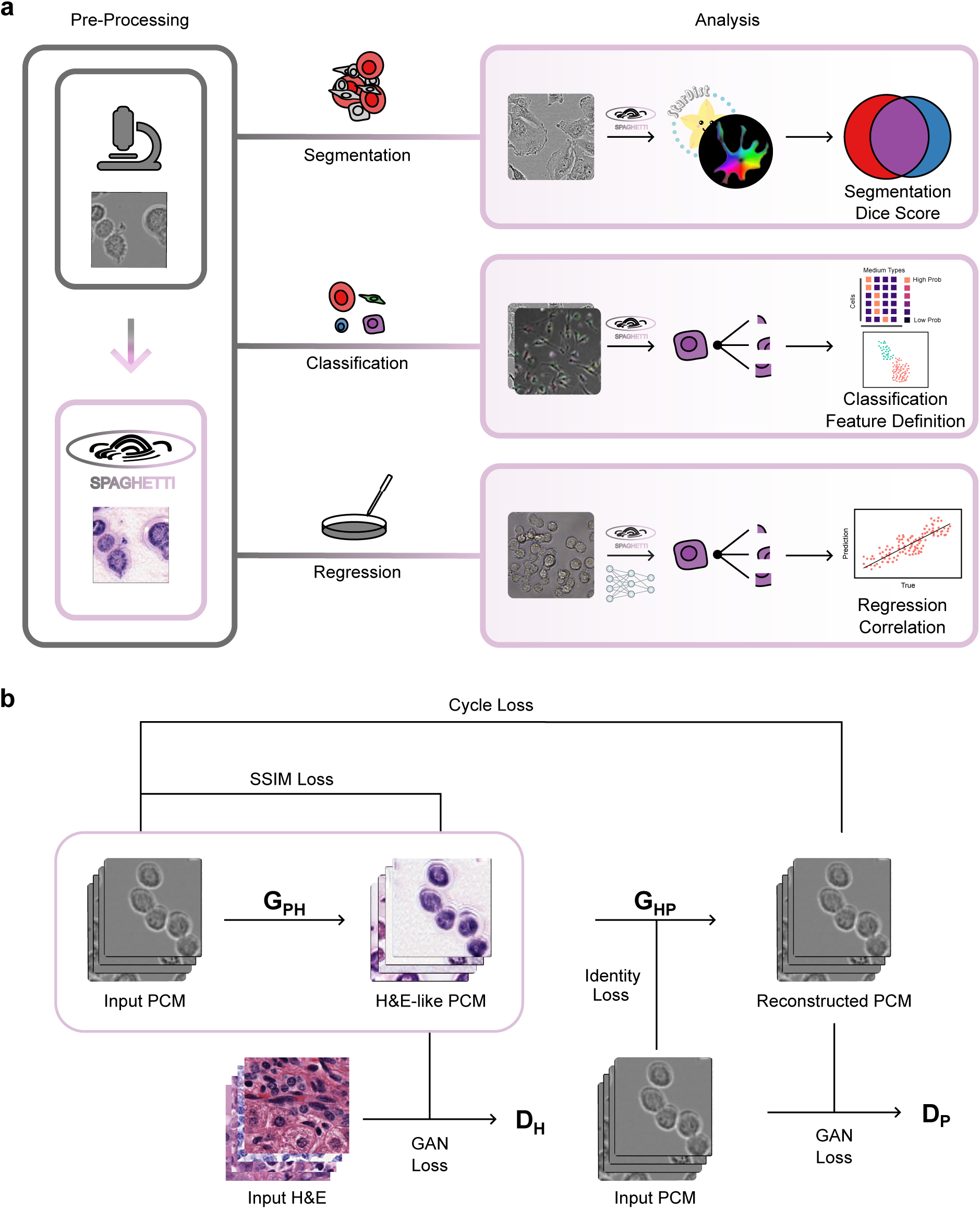
Overview of SPAGHETTI. **a**, Overview of SPAGHETTI workflow examples in the quantitative analysis of PCM. SPAGHETTI is a pre-processing step before running massive H&E models on PCM images for use with downstream tasks such as segmentation, cell type and media classification, and cell viability regression. **b**, The training and inference architecture of SPAGHETTI. SPAGHETTI learns to translate PCM images to H&E-like images using a cycle-consistent GAN architecture with an additional SSIM loss function. During training, a PCM to H&E generator (*G_PH_*) translates PCM images to H&E-like images, while the H&E to PCM generator (*G_HP_*) translates H&E images back to PCM-like images. The discriminators *D_P_* and *D_H_* distinguish between real and fake images for PCM and H&E images, respectively. The final SPAGHETTI model uses *G_PH_* to translate PCM to H&E-like images for use with downstream H&E models.

## Results

### SPAGHETTI introduces SSIM loss to adapt H&E tissue morphological models to PCM images

We present SPAGHETTI, a new architecture with a CycleGAN foundation, which improves downstream segmentation and classification tasks for PCM data through conversion of PCM images to H&E-like images. SPAGHETTI consists of two generators and two discriminators. We train the generators, which use a ResNet backbone^19^, to translate PCM images to H&E-like images and vice versa, while the discriminators, which use a PatchGAN architecture^20^, learn to distinguish between real and fake images produced by the generators. We optimize the objective function by solving a min-max problem, where generators minimize the loss function, while discriminators maximize the loss function. Once the training is completed, SPAGHETTI uses the PCM to H&E generator to convert PCM images to H&E-like images. However, this CycleGAN-based architecture hallucinates features in the translated images, where generated images contain features that are not present in the input images due to an unbalanced training dataset^21^. To address this issue, we introduce a new Structure Similarity Index Measurement (SSIM) loss. SSIM loss penalizes the generator if the translated images are not consistent with the input images by introducing or neglecting additional features, resulting in images that resemble key features of H&E images with less hallucination such as phantom nuclei compared to the original CycleGAN architecture (Figure 1b). Instead of visually inspecting these images, we use SPAGHETTI outputs as a preprocessing step for large tissue and H&E cell segmentors (e.g. Cellpose^7^ and StarDist^8^) as well as H&E feature extractors (e.g. Phikon-v2^10^, H-Optimus-0^9^, and UNI^11^), effectively adapting these commonly-used models for PCM data.

### SPAGHETTI assists H&E tissue and cell segmentation models in segmenting single cells

To demonstrate the utility of SPAGHETTI’s translation as an important preprocessing step for various H&E-based models, we first benchmarked the model’s segmentation performance. Cell segmentation is a critical step in nearly every biological image analysis pipeline. Yet, unlike H&E and other stained images, PCM images are harder to segment as the unstained cell boundaries are often not as clear as their stained counterparts. Therefore, despite the existence of many sophisticated models for segmenting H&E and other stained tissue images, few exist for PCM images. To determine the segmentation performance of SPAGHETTI, we first sought to compare segmentation outputs with the original CycleGAN architecture with SPAGHETTI-translated images using a dataset consisting of time-lapse PCM images of eight different cell types, displaying various morphological characteristics^22^. The introduction of SSIM loss by SPAGHETTI greatly reduced hallucinations of phantom nuclei (Figure 2a). Confirming the improved accuracy of SPAGHETTI, we next bench-marked segmentation performance of PCM images of eight different cell lines at different confluencies^22^ using two popular cell segmentors, Cellpose^7^ and StarDist^8^, along with preprocessed versions with SPAGHETTI and CycleGAN. SPAGHETTI clearly improved the segmentation of both Cellpose and StarDist, detecting more cells with fewer incorrect segmentations (Figure 2b). Quantifying the overlap of ground-truth segmentations with predicted boundaries with Dice scores, we found that SPAGHETTI significantly improved Cellpose’s tissuenet_cp3 model by 53.3 % (two-tailed paired T-test: *p* < 2.20 × 10^−16^) and StarDist’s 2D_versatile_he model by 95.0 % (two-tailed paired T-test: *p* < 2.20 × 10^−16^; Figure 2c). The performance of all of the scores were higher than the baseline segmentation result that used raw PCM images with the cyto3 model, a general cell segmentor commonly used in segmenting PCM images. Importantly, SPAGHETTI outperformed the original CycleGAN architecture for translation-based segmentation, showing the importance of the SSIM architecture (two-tailed paired T-test: *p* < 2.20 × 10^−16^; Figure 2c). These findings suggest that SPAGHETTI improves the performance of cell segmentors on PCM images, which enables more in-depth analyses of single-cell states and cell tracing experiments.

**Figure 2:**
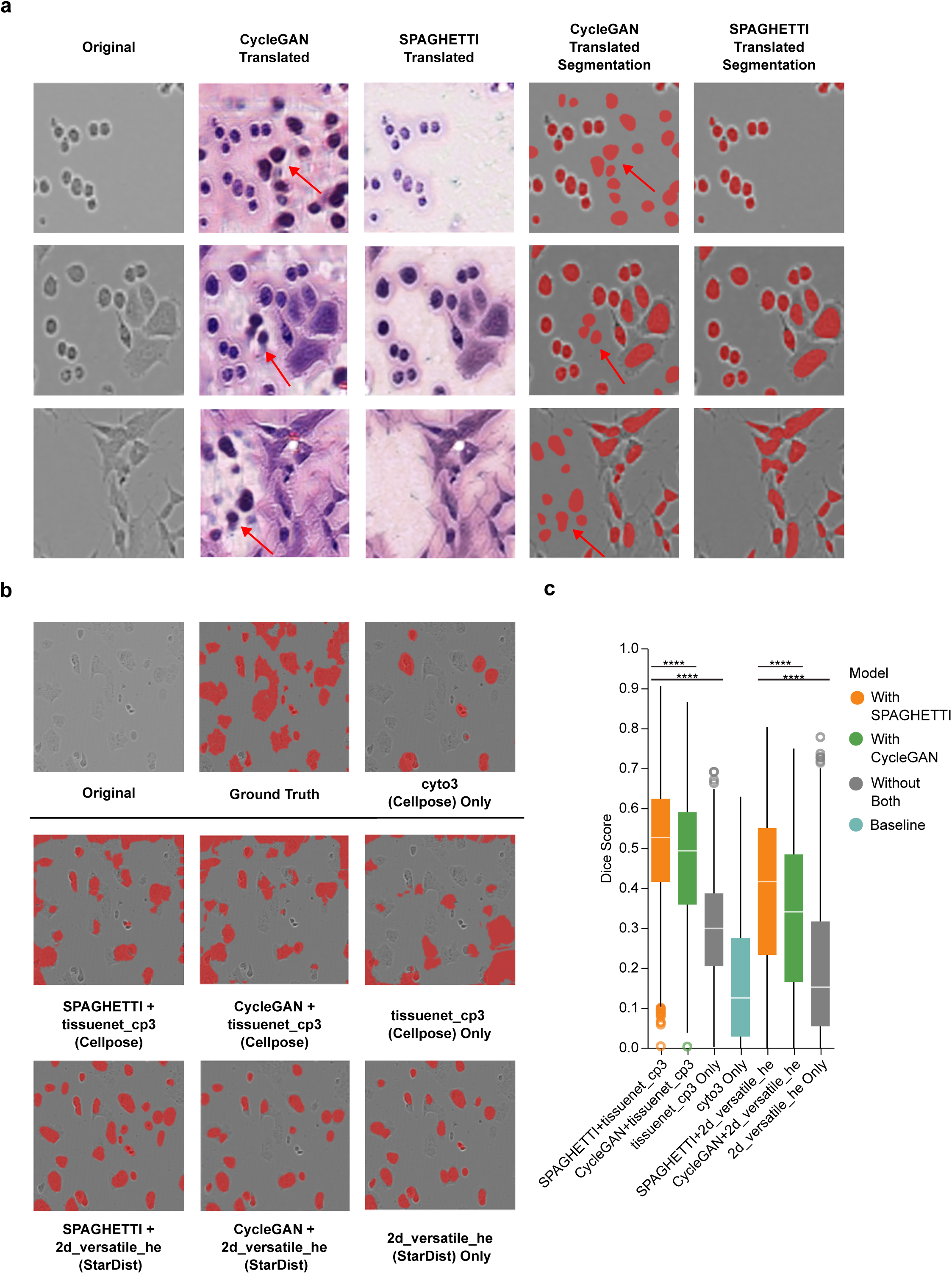
SPAGHETTI generates fewer hallucinations and improves cell segmentation with existing tissue and H&E models. **a**, Examples of original, CycleGAN-, and SPAGHETTI-translated images (first three columns left to right, respectively) and their cell segmentation results from the LIVECell dataset^22^ (right two columns). Red arrows indicate hallucinations from CycleGAN that do not appear in the original and are missing in SPAGHETTI-translated images. This reduction in hallucinations resulted in fewer false positives in the cell seg-mentation with SPAGHETTI-translated images. **b**, Examples of the cell segmentation results from Cellpose and StarDist on the SPAGHETTI-, CycleGAN-translated, and original PCM images from the LIVECell dataset. **c**, Box-and-whisker plots (center line, median; box limits, upper (75^th^) and lower (25^th^) percentiles; whiskers, 1.5 × interquartile range; points, outliers) of Dice scores reporting cell segmentation performance from Cellpose and StarDist on the SPAGHETTI-or CycleGAN-translated and original PCM images from the LIVECell dataset. Preprocessing with SPAGHETTI significantly improved the performance of all tested models. Two-tailed paired T-test: *p* < 2.20 × 10^−16^.

### SPAGHETTI improves performance of cell-type annotation

Cell type classification typically follows most segmentation tasks to better characterize and track cell abundance^23^. We determined SPAGHETTI’s impact on established cell-type annotators by collecting PCM images from eight different cell lines taken across several days at different confluencies^22^. We applied UNI, one of the largest state-of-the-art (SOTA) Vision Transformer (ViT) based foundational feature extractors designed for H&E^11^ based on trained dataset size, H-Optimus-0, one of the largest SOTA ViT feature extractors based on the number of model parameters for H&E^9^, and Phikon-v2, one of the newest SOTA ViT feature extractor model for H&E^10^, coupled with a random forest cell-type classifier with and without SPAGHETTI. In all cases, SPAGHETTI in addition to each model significantly increased performance (two-tailed permutation test: *p* = 0.0171, *p* < 1.00 × 10^−5^, and *p* = 0.0450 respectively for Phikon-v2, H-Optimus-0, and UNI; Figure 3a and Supplementary Figure S1).

**Figure 3:**
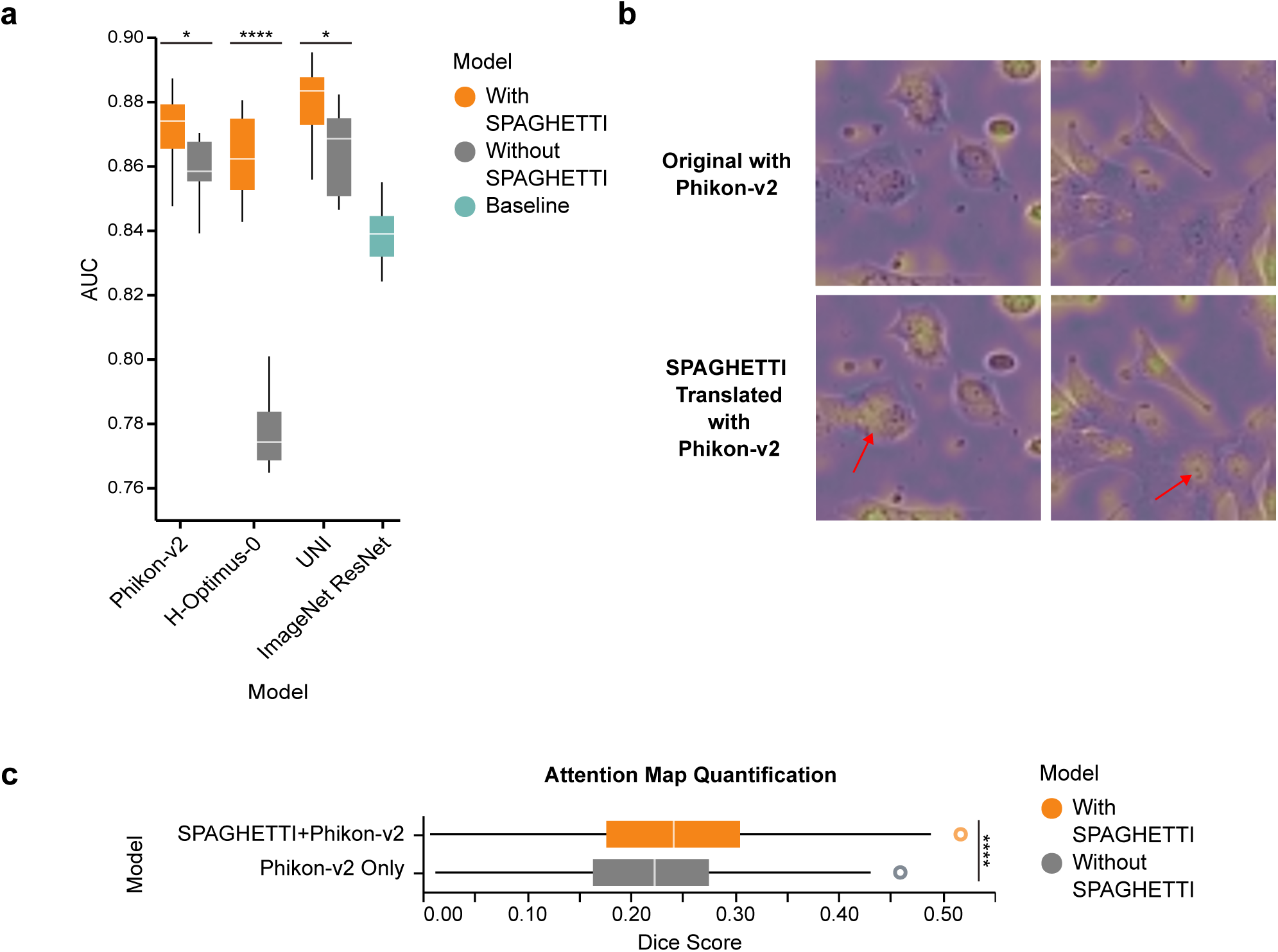
SPAGHETTI improves cell-type annotation performance and focuses feature attention of H&E feature extractors on the LIVECell dataset^22^. **a**, The AUC values from micro receiver operating characteristic (ROC) curves for LIVECell cell-type annotation using a random forest classifier on features extracted using Phikon-v2, H-Optimus-0, and UNI from SPAGHETTI-translated PCM or original PCM images. SPAGHETTI-translated images significantly improved the performance of all classifiers compared to raw PCM images. **b**, Attention maps examples of the *CLS* token from Phikon-v2 on the SPAGHETTI-translated and original PCM images, where the brighter colour indicates the higher attention score. The SPAGHETTI-translated images allow the model to focus better on cell populations (indicated with red arrows). **c**, Box-and-whisker plots of probabilistic Dice scores evaluating overlap of attention masks generated by Phikon-v2 on the SPAGHETTI-translated and original images with segmentations from the LIVECell dataset. Probabilistic Dice scores are significantly higher when using SPAGHETTI-translated images compared to using the original PCM images. *: *p* < 0.05, ****: *p* < 0.0001

Interestingly, while UNI had the best classification performance when used alone at 0.870 AUC, the two overall highest performing methods were SPAGHETTI with UNI at 0.882 AUC and SPAGHETTI with UNI at 0.875 AUC, demonstrating significant gains when using SPAGHETTI on much smaller models making them more generalizable when applied to different imaging modalities. This gain is important, as while using raw PCM images with Phikon-v2 and UNI outperformed the pre-trained ImageNet ResNet model at 0.840 AUC which is often used as a general feature extractor for medical images^24^, using raw PCM images with H-Optimus-0 at 0.770 AUC failed to outperform this baseline. However, using SPAGHETTI-translated images can outperform this baseline in all tested feature extractors.

To understand potential factors contributing the success of SPAGHETTI, we investigated the attention maps of the feature extractor on the original and SPAGHETTI-translated images as part of this classification task. Interestingly, we found that SPAGHETTI with Phikon-v2 had significantly higher overlap of attention with cell location versus Phikon-v2 only, suggesting SPAGHETTI helps H&E feature extractors focus more on the cells themselves (two-tailed paired *t*-test, *p* < 2.20 × 10^−16^; Figures 3b and 3c). Although the Dice score is relatively low, this outcome is expected given that the experiment is not focused on segmentation, but rather aims to demonstrate a more concentrated allocation of attention on the cell. Together, these results suggest that SPAGHETTI can help existing ViT based feature extractors such as Phikon-v2 to better identify cell populations in the images, leading to well-defined features.

### Performance of media conditions prediction on an external dataset increases with SPAGHETTI-translated images

Although the model performs well with cell-type classification, morphological differences between cell types are relatively large compared to differences between cells of the same type. As such, we then benchmarked SPAGHETTI’s performance on media condition classification to demonstrate its utility and generalizability in classification on an external and independent dataset with cells that show more similar morphologies. Media condition greatly affects cell state and function. As such, predicting culture media from cell morphology can provide insights to the effect of different media on cells^25^ and can even be used to quantify drug responses in a clinical setting^26^. To ascertain the capability of SPAGHETTI to assist models in prediction culture media from cell morphology, we analyzed C2C12 cells from time-lapse PCM images of four media conditions containing different growth factors^27^. Interesting, as with cell-type classification, attention maps with SPAGHETTI-translated images followed by Phikon-v2 focused on cells rather than unspecific regions of each image (Figure 4a). Using a random forest classifier on cells across different media using the morphological features extracted by Phikon-v2, H-Optimus-0, and UNI, we saw overall improvement of area under the curve (AUC) values when using SPAGHETTI-translated images compared to raw PCM images for all tested feature extractors (two-tailed permutation test: all *p* < 1.00 × 10^−5^; Figure 4b and Supplementary Figure S2). These improvements were as high as 18.1 % for H-Optimus-0, increasing the performance to 0.910 AUC which is higher than the best performing model without SPAGHETTI, Phikon-v2, at 0.850 AUC.

**Figure 4:**
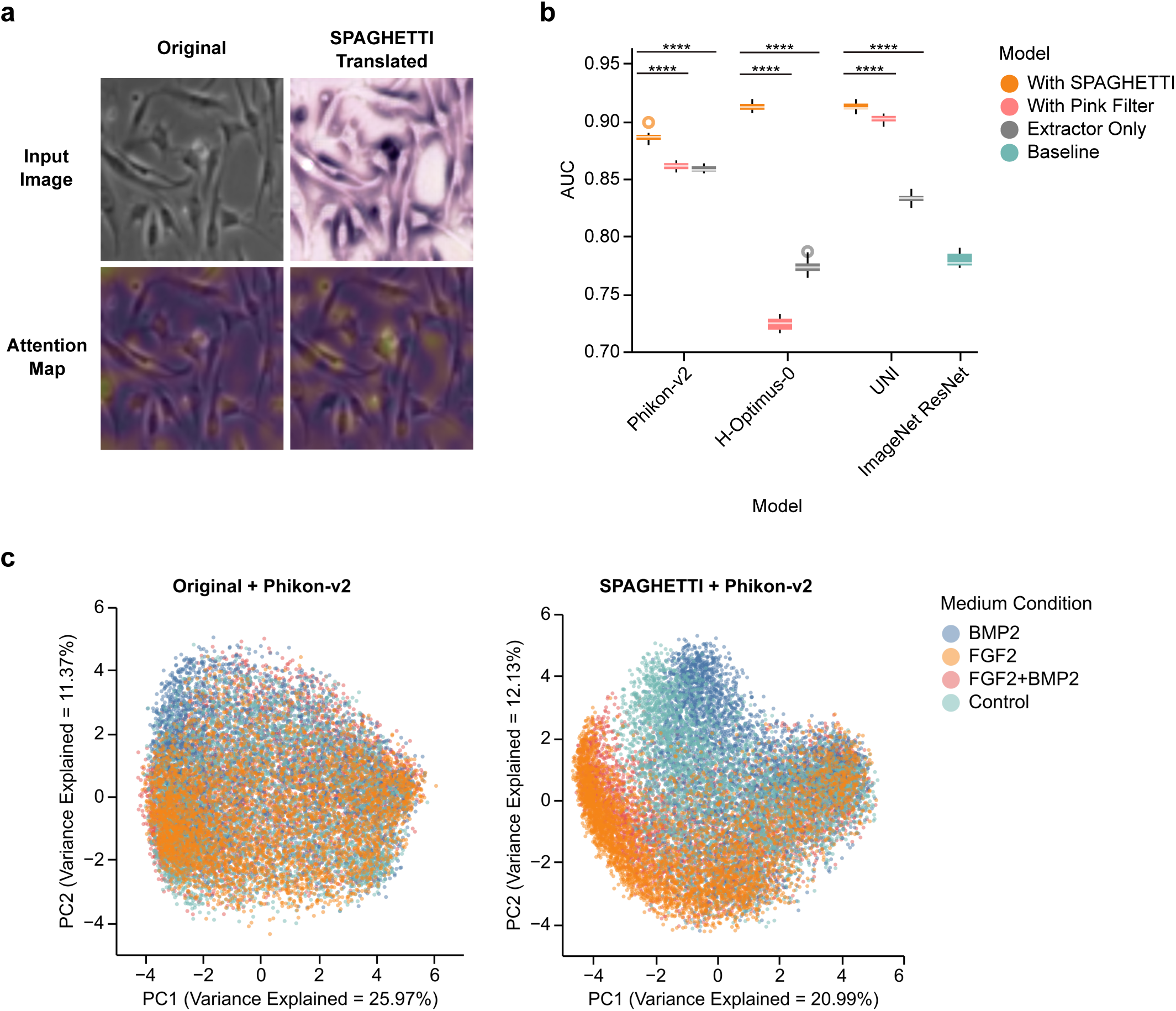
H&E translation from PCM images increases performance of media condition classification and produces better defined features in an external dataset of C2C12 cells in different media conditions^27^. **a**, Attention map examples of the *CLS* token from Phikon-v2 on the SPAGHETTI-translated and original PCM images from the C2C12 dataset, where brighter colours indicate higher attention score. The SPAGHETTI-translated images boost the model’s focus on cell populations. **b**, The micro ROC curves for classification of C2C12 media conditions with a random forest classifier on features extracted using Phikon-v2, H-Optimus-0, and UNI from SPAGHETTI-converted PCM, pink filter-transformed PCM, and original PCM images. SPAGHETTI-translated images improved performance over both the original and pink filter images. **c**, The principal component analysis (PCA) showing the first two principal components of the features extracted by Phikon-v2 from the SPAGHETTI-converted and original PCM images. The SPAGHETTI-converted features show a clearer separation between different media treatments. ****: *p* < 0.0001

To investigate if SPAGHETTI’s translated images learned the underlying structure of cells in this independent dataset, we compared SPAGHETTI with a trivial pink filter transformation (Supplementary Figure S3). In all cases, SPAGHETTI indeed displayed major improvements over the pink filter, confirming SPAGHETTI’s capability of producing helpful H&E-like images (two-tailed permutation test, all *p* < 1 × 10^−6^). Surprisingly, we observed that the pink filter transformation can either positively or negatively affect media prediction depending on the downstream feature extractor. We observed that the pink filter transformation had the smallest effect on performance when paired with Phikon-v2, the model trained on the largest dataset (Supplementary Table S1). Together with the observation that Phikon-v2 outperformed over other non-translated images, this result suggests that larger training datasets can increase robustness to colour transformations and generalizability to focus on key features of images.

To see whether the features identified by massive models such as Phikon-v2 led to distinct profiles for each tile, we used principal component analysis (PCA) to visualize SPAGHETTI-translated Phikon-v2 and the raw PCM Phikon-v2 features. We found a clear separation between the different experimental media in the SPAGHETTI-translated features compared to the raw PCM features (Figure 4c). Together, our results suggest that SPAGHETTI can help improve the performance of H&E feature extractors on various prediction tasks by not only directing model attention towards cells, but also leading to better defined feature extraction.

### Self-supervised retraining enables SPAGHETTI to predict cell viability on stained PCM datasets

To determine whether SPAGHETTI can facilitate regression-based analyses on cell morphology changed by staining, we applied SPAGHETTI to predict cell viability of human colon adenocarcinoma Caco-2 cells^28^. These cells were treated with topoisomerase inhibitor camptothecin and stained with either Acridine Orange or DioC6 and propidium iodide, resulting in green and red channels to measure cell viability. Cell viability is a key measure in drug development to determine and compare the efficacy of different treatments. Yet to measure cell viability, researchers often rely on staining methods and assays for quantitative measurements, which are expensive and time-consuming^29^. To determine whether the SPAGHETTI architecture could generalize to stained cells, we applied our model to assist a linear regression model to predict cell viability from PCM images using the features obtained from translated images through a retraining process (Figure 5a).

**Figure 5:**
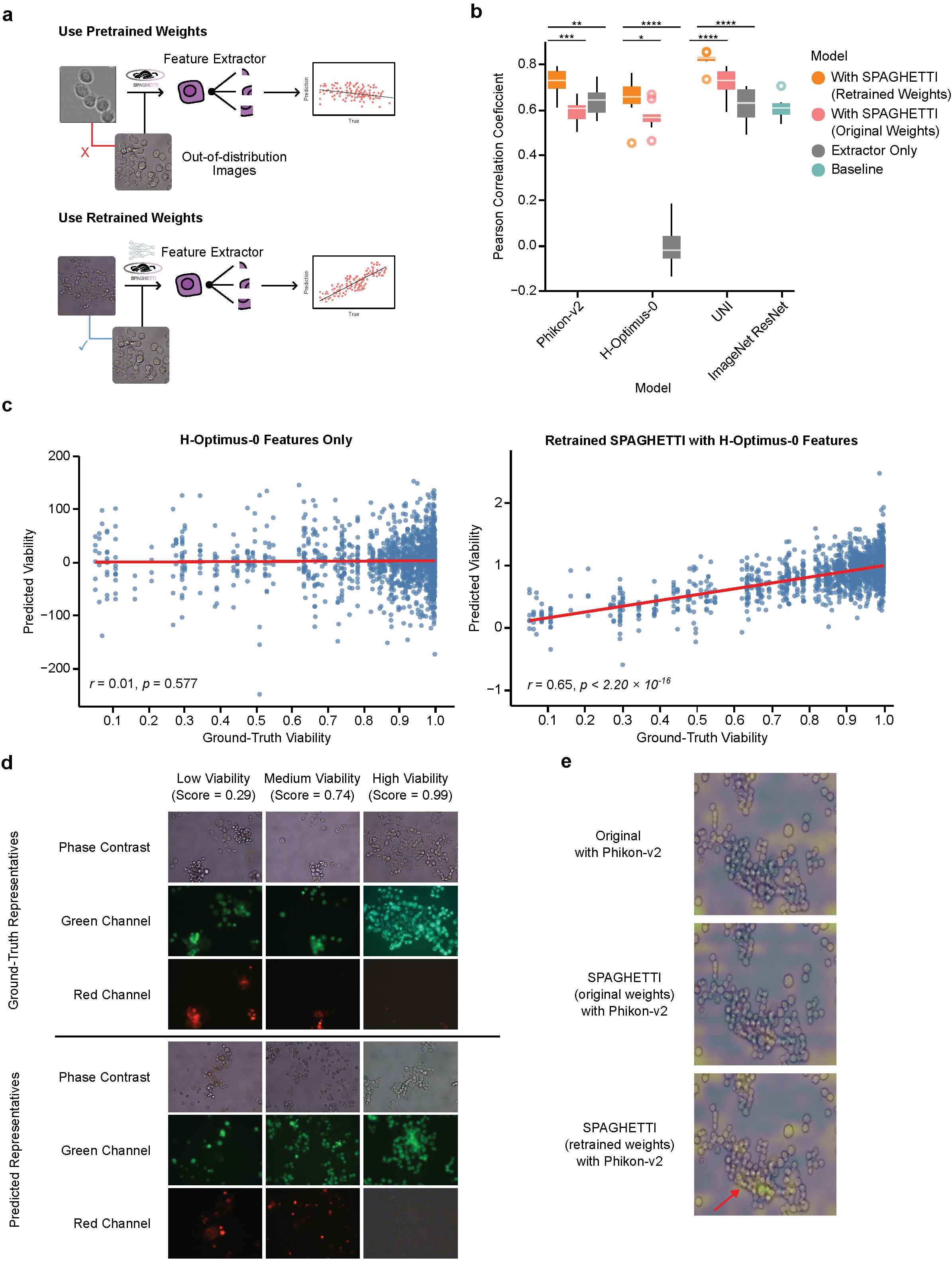
Self-supervised learning improves SPAGHETTI performance on an out-of-distribution PCM dataset of human colon adenocarcinoma Caco-2 cells^28^. **a**, Workflow comparison showing the original SPAGHETTI weights (top) and retraining SPAGHETTI on the Caco-2 dataset^28^ (bottom). Retraining helps with image translation for stained cells. **b**, Pearson correlation coefficients between the predicted and ground-truth viability scores across different feature extractors. Features from the retrained SPAGHETTI model significantly improved the correlation compared to those from the original SPAGHETTI model and the original PCM image. **c**, Scatter plot of the predicted (*y* axis) and ground truth (*x* axis) viability scores from the linear regression on features from the original PCM images (left) or from translated images using retrained SPAGHETTI (right) using H-Optimus-0. The features from the retrained SPAGHETTI model are much better correlated with the ground-truth viability scores. **d**, Examples of raw PCM images (top row) and their corresponding green and red channel images (bottom two rows) representing low, medium, and high viability scores. The increasing levels from the green channel and decreasing levels of red channel correspond to increasing viability scores at similar levels for both the ground-truth scores (top three rows) and predicted scores (bottom three rows). **e**, Attention map examples of the *CLS* token from Phikon-v2 on the retrained SPAGHETTI-translated, original SPAGHETTI translated, and original PCM images. The retrained SPAGHETTI-translated images allow the model to focus better on cell populations (red arrow). *: *p* < 0.05, **: *p* < 0.01, ***: *p* < 0.001, ****: *p* < 0.0001.

Although we previously applied SPAGHETTI to PCM images, the process of staining cells may cause morphological variations. As such, there were clear differences in images from stained Caco-2 cells distinct from PCM images from the LIVECell dataset previously used to train SPAGHETTI. These out-of-distribution PCM images resulted in lower predicted accuracy of cell viability for Phikon-v2, likely due to the translated images having less resemblance to H&E images (Figure 5b and Supplementary Figure S4). To overcome this obstacle and generalize the use of SPAGHETTI in the case of out-of-distribution data, we propose a workflow that involves the self-supervised retraining of the SPAGHETTI architecture. This workflow significantly improved the correlation and reduced errors between the predicted and actual cell viability for all feature extractors paired with the regression compared to the untranslated images (two-tailed permutation test: *p* = 9.34 × 10^−3^ for Phikon-v2, and *p* < 1.00 × 10^−5^ for both H-Optimus-0 and UNI) and the original SPAGHETTI weights (two-tailed permutation test: *p* = 2.62 × 10^−3^, *p* = 0.0219, and *p* = 2.43 × 10^−4^ respectively for Phikon-v2, H-Optimus-0, and UNI; Figure 5b and Supplementary Figure S5). The results were drastically improved with H-Optimus-0, whose correlation improved overall by 0.7 using retrained SPAGHETTI compared to using H-Optimus-0 directly on the PCM images, resulting in a clear linear relationship between the predicted and ground truth cell viability scores (Figure 5c).

For additional verification of our predicted viability scores, we manually inspected images at the low, medium, and high viability scores. From low to high viability, we observed a monotonic increase in the green channel and decrease in the red channel at similar levels when comparing images at each score for predicted and training datasets (Figure 5d). We reasoned that this alignment with the ground truth may relate to higher overlap of attention on the cell population (Figure 5e). As the retrained SPAGHETTI outperformed the original SPAGHETTI weights and the non-translated PCM for all feature extractors, we propose that retraining SPAGHETTI on a new PCM dataset can help adapt the model and improve the performance of H&E feature extractors for that dataset. Retraining SPAGHETTI is much less computationally expensive compared to retraining a feature extractor, as SPAGHETTI only requires a few hundred PCM images to train and converges quickly despite the small training dataset size (Supplementary Figure S6 and Supplementary Table S1). Coupled with the self-supervised training of SPAGHETTI, retraining is a straightforward process using a small subset of the PCM images in the target dataset and a publicly available H&E dataset such as the PanNuke dataset^30^ using the training function provided in the SPAGHETTI package. This new workflow enables the use of SPAGHETTI on a wide variety of PCM datasets to better assist H&E feature extractors in downstream analyses.

### Smaller models outperform larger models by using SPAGHETTI-translated images

Despite the significant performance improvements SPAGHETTI brings to existing models, the method is relatively fast and computationally inexpensive. SPAGHETTI with feature extractors requires only ≈ 100 MB more GPU memory and ≈ 0.140 s more runtime per image compared to running the feature extractors alone (Figures 6a and 6b). To determine whether these performance gains were offset by smaller training dataset sizes, we ran-domly drew subsets of data from LIVECell and C2C12 for each model with and without SPAGHETTI. Interestingly, models with SPAGHETTI still outperformed models without preprocessing, even down to 10% of the complete dataset (Figure 6c). This robustness suggests that SPAGHETTI can improve the generalizability of the feature extractors.

**Figure 6:**
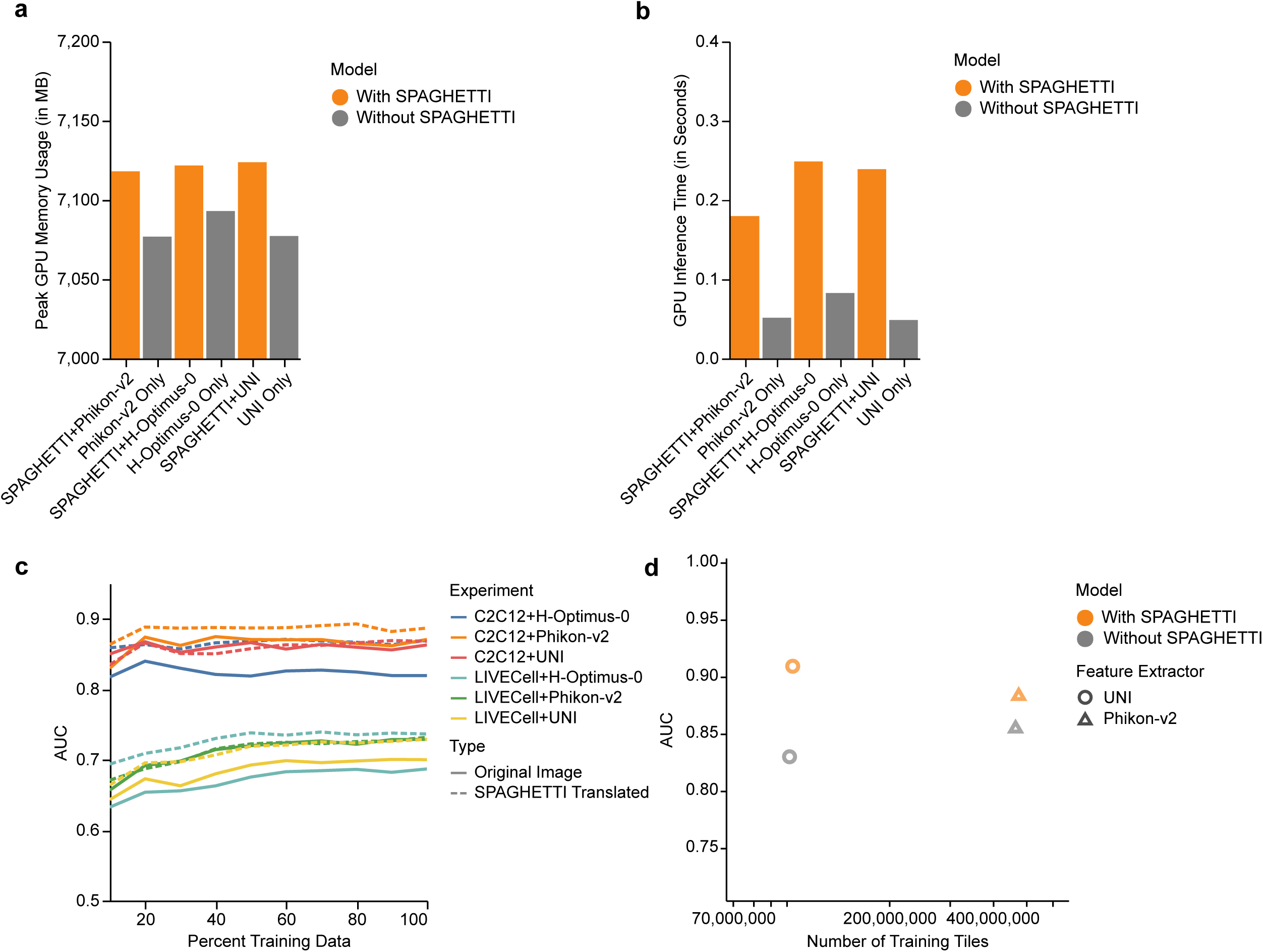
SPAGHETTI enables lightweight models to outperform heavier models with little overhead. **a**, Bar plot of the memory allocated to run SPAGHETTI with different feature extractors on the C2C12 dataset^27^. SPAGHETTI only requires ≈ 100 MB more GPU memory compared to feature extractors alone, which require over 7 GB of GPU memory allocation. **b**, Bar plot of the runtime for SPAGHETTI and feature extractors on the C2C12 dataset. SPAGHETTI only requires milliseconds additional time per image. **c**, Micro ROC curves for classification of LIVECell cell types and C2C12 media treatments, using a random forest classifier on features extracted using Phikon-v2, H-Optimus-0, and UNI from SPAGHETTI-converted PCM and original PCM images, across training dataset sizes. In all cases, SPAGHETTI increased performance. **d**, Training dataset sizes and AUC values from the micro ROC curves for classification of C2C12 media conditions across all models with and without SPAGHETTI preprocessing. While Phikon-v2 outperformed UNI without SPAGHETTI, the combination of UNI and SPAGHETTI can outperform Phikon-v2 alone, which together use hundreds of millions less data points for training.

SPAGHETTI is a lightweight model as its generator has a convolutional-neural-network-based ResNet architecture backbone. Compared to transformer-based foundational feature extractors, SPAGHETTI’s architecture greatly reduces model parameter numbers, reducing the dataset size required for training and the computational resources required to run. As such, while transformer-based foundational feature extractors are often trained on millions if not billions of images, we trained SPAGHETTI with only a few thousand unpaired images (Supplementary Table S1). We inquired whether SPAGHETTI with another lightweight model could bring the overall performance in line with a massive model in the media classification task. Surprisingly, not only did SPAGHETTI increase the performance of UNI, the model with the smaller training dataset, but improved the model to such an extent that it outperformed Phikon-v2, a model requiring hundreds of millions more tiles to train (Figure 6d). This result suggests that SPAGHETTI can deliver a synergy with other feature extractors to improve the performance of the models without requiring a large amount of data to train. As such, incorporating SPAGHETTI into an analysis pipeline not only improves the generalizability of downstream analyses but also is computationally cheap and can easily integrate into existing workflows through our PyPI package or the Docker image.

## Discussion

PCM, despite having great power in capturing cell dynamics in temporal experiments, bypassing staining while also providing informative measurements of cell morphology, are underutilized in quantitative analysis. This untapped potential may be due to few algorithms working with PCM images caused by the lack of large training datasets. Here, we proposed a solution to this problem by introducing SPAGHETTI, an image translator using a cycle-consistent generative adversarial network and an additional SSIM loss function.

We measured the capability of SPAGHETTI to translate PCM images into images re-sembling H&E pathological images and saw that the SSIM loss function greatly reduced the amount of hallucinations in resulting images. We showed that SPAGHETTI can help state-of-the-art massive H&E feature extractors better identify cell populations in images, leading to more relevant feature attention as a result. We demonstrated that using SPAGHETTI as a preprocessing step improved performance on a variety of downstream tasks such as cell segmentation, cell-type annotation, experimental media classification, and cell viability prediction across several independent datasets using a variety of H&E based models. Importantly, we demonstrate how SPAGHETTI can improve generalizability and assist lightweight models trained on smaller datasets to outperform models trained on larger datasets.

Although SPAGHETTI enables the use of sophisticated segmentors and feature extrac-tors on PCM data, there are some limitations. While we demonstrated that SPAGHETTI can improve the performance of H&E morphological models on PCM images, the trans-lated images may not be biologically accurate as if cells were actually stained with H&E. For example, typical H&E images are taken from sections that may expose the inside of cells, while our tested PCM images do not have such sections. This lack of sectioning motivates SPAGHETTI to be used specifically as a preprocessing step for downstream analyses instead of being used for direct visual examination. Future work could involve the development of a more accurate model that can generate biologically accurate H&E-like images from PCM images so that translated images may pass direct visual examination. In addition, the SPAGHETTI architecture may inspire future models looking at other imaging modalities such as differential interference contrast microscopy and bright field microscopy to further leverage large models for popular and useful modalities that lack big training datasets. Furthermore, while retraining SPAGHETTI is a relatively inexpensive process, a GPU is still necessary. Future versions could involve a client-server model that enables the retraining online using a few hundred PCM images and a publicly available H&E dataset, making the retraining process even more accessible. As models designed for H&E leverage morphological features to infer cell state, SPAGHETTI provides a novel way to transfer those abilities to PCM images, enabling new discoveries through the temporal monitoring and analyses of cells.

## Methods

### SPAGHETTI model architecture

SPAGHETTI is built upon the original CycleGAN architecture, which uses a generative adversarial network (GAN) for image translation^16^. Given a set of images from PCM (*P*) and H&E (*H*), the objective of the model is to learn two mappings, *G_PH_*: *P* → *H* and *G_HP_*: *H* → *P*, between a training set of *n* PCM images (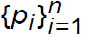, where *p_i_* ∈ *P*) and *m* H&E images (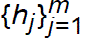, where *h_j_ i*=1 ∈ *H*). Importantly, *m* and *n* do not need to be equal, nor do images need to be paired (i.e. *p_i_* and *h_j_* may be from different samples). To learn these mappings, we employed two adversarial discriminators, *D_P_* and *D_H_*, to distinguish between real and fake images generated by *G_PH_* and *G_HP_*.

SPAGHETTI uses four different loss functions. Three are found in the original CycleGAN architecture^16^, including adversarial loss (ℒ*_GAN_* for both *G_PH_* and *G_HP_*), cycle-consistency loss (ℒ*_cyc_*), and identity loss (ℒ*_idt_*). In addition, SPAGHETTI also employs a new SSIM loss (ℒ*_SSIM_*).

The adversarial loss helps train both generators to produce realistic images by ensuring that the discriminators cannot distinguish between real and fake images produced by the generators. We define adversarial loss using the binary cross entropy loss function:

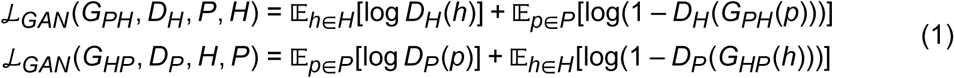

We use cycle-consistency loss to ensure that the regenerated images are consistent with the input images. We define cycle-consistency loss by the *L*_1_ norm of the difference between the input and the regenerated images for both domains:

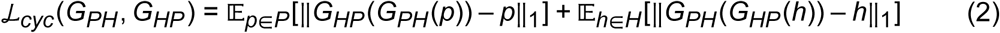

The identity loss ensures that the generators do not alter the input images if they are passed through the generator from the same domain. This loss can preserve the colour and texture of the input images when passed through the translation process. We define identity loss by the *L*_1_ norm of the difference between the input and the generated images for both:

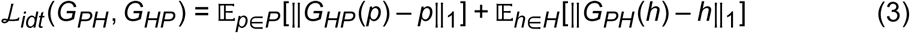

We derive the SSIM loss from the SSIM value, which we determine with a sliding window and averaged across the entire image. The SSIM value is a measure of the similarity between two images based on luminance, contrast, and structure, which ranges from –1 to 1, where 1 indicates that the two images are identical^31^. Given two images *x* and *y*,

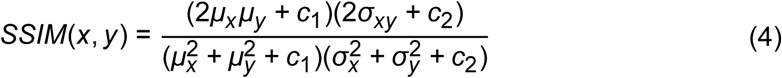

where *μ_x_* and *μ_y_* are the average of *x* and *y*, 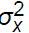 and 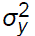 are the variance of *x* and *y*, *σ_xy_* is the covariance of *x* and *y*, and *c*_1_ and *c*_2_ are constants to stabilize the division with weak denominator. Since the SSIM value ranges from –1 to 1 and measures the similarity between two images, we then define SSIM loss as the difference between 1 and the SSIM value so that the loss is minimized when the two images are identical and have an SSIM value of 1. We then average the SSIM loss across all images for both domains of *P* and *H*:

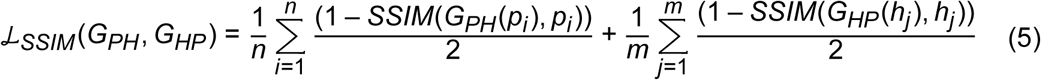

The overall loss is a combination of all the above losses, which is defined as follows:

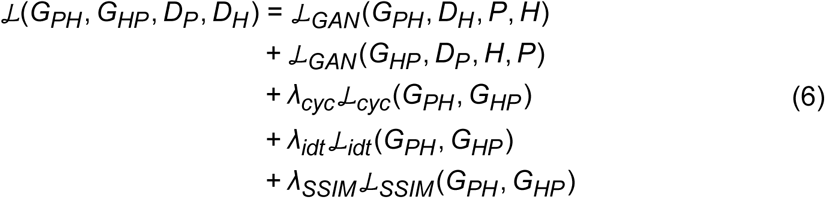

where *λ_cyc_*, *λ_idt_*, and *λ_SSIM_* are the hyperparameters that control the relative importance of the cycle-consistency, identity, and SSIM loss terms, respectively. The overall objective function is optimized by solving the following min-max problem, where the generators are optimized to minimize the loss function, while the discriminators are optimized to maximize the loss function:

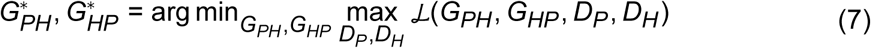

### Training details

We trained SPAGHETTI and CycleGAN architectures using the LIVECell dataset^22^ for PCM (*n* = 5,326) and the PanNuke dataset^30^ for H&E (*m* = 7,904). We resized all images to 256×256 pixels and converted the range of the pixel values to [0, 1]. We normalized the pixel values to a mean of 0.5 and a standard deviation of 0.5, and applied random crops, flips, and rotations for data augmentation. We implemented generators using a ResNet architecture^19^ with 9 residual blocks. We implemented discriminators using a PatchGAN architecture^20^ with 16×16 patches. We split the dataset to 80% training and 20% testing, before using a batch size of 16 and the AdamW optimizer^32^ with a learning rate of 0.001 and a weight decay of 0.01 for training. We set hyperparameters *λ_cyc_* and *λ_idt_* to 10.0 and 5.0 for both SPAGHETTI and CycleGAN architectures, and *λ_SSIM_* to 10.0 for SPAGHETTI. We trained both models for 100 epochs on 8 NVIDIA T4 Tensor Core GPUs on the Vaughan High Performance Clusters at the Vector Institute for Artificial Intelligence. After training, we used the PCM to H&E generator 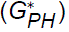 as the generators for SPAGHETTI and CycleGAN to generate H&E-like images from PCM images.

### Feature attention visualization and quantification

We visualized the attention map of the *CLS* token, which is used as the feature of image^33^, from the last transformer layer of Phikon-v2^10^ on the 20% testing portion of the LIVECell dataset, the C2C12 experimental media dataset^27^, and the Caco-2 dataset^28^. We reshaped the *CLS* token to be the shape of 14×14 pixels, before bicubic extrapolation back to the original image size of 256×256 pixels. We then computed the attention map by averaging the attention weights across all heads and normalizing the attention weights to the range of [0, 1] per image. Lastly, we overlaid the attention map on the original image to visualize feature attention by utilizing a plasma colour map.

To quantify the feature attention for LIVECell, we computed the probabilistic Dice-Sørensen coefficient (Dice score) between the attention map and the ground truth cell mask by treating the normalized attention map as a segmentation mask. The probabilistic Dice score is a measure of the overlap between a ground truth mask (*A*) and a predicted mask (*B*), where a higher Dice score indicates a better overlap between the two masks:

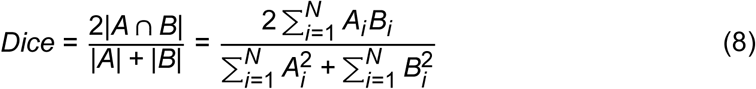

We repeated the above process for using SPAGHETTI as the preprocessing step and using no special preprocessing steps before inputting the images into Phikon-v2.

### LIVECell cell segmentation

We performed cell segmentation using two popular existing cell segmentors Cellpose^7^ and StarDist^8^ on the 20% testing portion of the LIVECell dataset. In particular, we used the tissuenet_cp3 model for Cellpose and the 2D_versatile_he model for StarDist. We reported the segmentation results using the binary Dice score, which uses the same formula as Equation (8) but now both the ground truth cell masks and predicted cell masks are binary masks of either 1 (cell) or 0 (background).

We repeated the above process for using SPAGHETTI as the preprocessing step and using the original PCM images before feeding images into cell segmentors. As a control for Cellpose, we also passed the original PCM images through the cyto3 model of Cellpose. We performed two-tailed paired *t*-tests to compare the significance of Dice scores improvements from using SPAGHETTI as the preprocessing step.

### Cell-type and experimental media classification using image features

We performed cell-type classifications on the testing dataset consisting of 20% of the total LIVECell images as well as media predictions on the external C2C12 dataset by training a random forest classifier with 100 trees and a maximum depth of 2. The LIVECell dataset contains eight different cells types at various confluencies. The C2C12 dataset consists of PCM images of C2C12 cells under different culture media conditions: control (no growth factors), fibroblast growth factor 2, bone morphogenetic protein 2, and a combination of fibroblast growth factor 2 and bone morphogenetic protein 2. We used Phikon-v2, H-Optimus-0^9^, and UNI^11^, three state-of-the-art foundational models for H&E slides benchmarked by external sources^34,35^, as feature extractors, as well as a ResNet model pre-trained on ImageNet as the baseline feature extractor. The task was to use the morphological features extracted from each model to classify the image into one of the eight cell types annotated by the LIVECell dataset, or one of the four media conditions annotated by the C2C12 dataset. In addition to image resizing and normalization to both LIVECell and C2C12 datasets, we added an additional colour inversion step to the C2C12 dataset to ensure that the background colour better resembles the training LIVECell PCM. We reported the AUC values from the ROC plots and their standard deviations of each random forest classifier using a 10-fold cross-validation approach. We repeated the above process using SPAGHETTI as the preprocessing step, and using the original PCM images before feeding the images into the feature extractors. For the C2C12 data, we also applied a pink filter (RGB values of [194, 128, 163]) by averaging the filter with the original image. We reported ROC-AUC values and used a permutation test of 100,000 times to determine significance.

### Varying training dataset size

To evaluate the generalizability of SPAGHETTI, we trained features extracted using various models on the testing set of LIVECell and C2C12 datasets for downstream classification tasks using different training dataset sizes. We divided the dataset with a ratio of 80% training and 20% testing images, and used a range of training dataset sizes from 10% to 100% of the original training dataset size. We repeated the above process using SPAGHETTI as the preprocessing step and the original PCM images before feeding the images into each feature extractor. We reported the AUC values from the ROC plots of the random forest classifiers on the testing set.

### Cell viability prediction and retraining

We used the Caco-2 dataset^28^ to predict cell viability. The Caco-2 dataset contains cells that are both non-treated and treated with camptothecin, a topoisomerase inhibitor, and stained with propidium iodide and Acridine Orange or DioC6. We calculated the cell viability score by using the number of pixels greater than 60 in the red channel (stained with propidium iodide) divided by the either the number of pixel greater than 60 in the green channel (stained with Acridine Orange) or the total number of pixels greater than 60 in both the red and green channels (stained with DioC6). For each field of view, a PCM image, a red channel image, and a green channel image are provided. We retrained the model using 50% of the dataset and use the other 50% for validation by following the training procedure in *Training details*. We then performed linear regression on the features extracted from the retrained SPAGHETTI translated images, original SPAGHETTI translated images, and the original PCM images using Phikon-v2, H-Optimus-0, UNI, and ImageNet ResNet as feature extractors. We reported the Pearson correlation coefficient between the predicted and actual viability scores from a 10-fold cross-validation approach, and computed the significance using a two-tailed permutation test of 100,000 times.

### Resource usage measurements

We measured the maximum GPU memory usage and the time taken to run SPAGHETTI and the feature extractors on the C2C12 dataset on an NVIDIA T4 Tensor Core GPU on the Vaughan High Performance Clusters at the Vector Institute for Artificial Intelligence by using the torch.cuda.max_memory_allocated() function from PyTorch and the time module in Python 3.9.

## Data Availability

The LIVECell dataset used for training and validation can be obtained at http://livecell-dataset.s3.eu-central-1.amazonaws.com/LIVECell_dataset_2021/images.zip. The PanNuke dataset used for training can be obtained at https://warwick.ac.uk/fac/cross_fac/tia/data/pannuke. The C2C12 dataset used for external validation can be obtained at https://osf.io/ysaq2/. The Caco-2 dataset used for cell viability prediction can be obtained at https://figshare.com/articles/dataset/VirtualStaining_Dataset/21971558/1?file=38982755.

## Code availability

SPAGHETTI is available at https://github.com/schwartzlab-methods/spaghetti or as a Docker image at https://hub.docker.com/repository/docker/yinnikun/spag hetti/. SPAGHETTI is also available on PyPI https://pypi.org/project/pcm-spagh etti/. Tutorials and example vignettes for using and training SPAGHETTI are available at https://github.com/schwartzlab-methods/spaghetti/tree/main/tutorials. Scripts for generating the figures and plots of the manuscript can be found at https://github.com/schwartzlab-methods/spaghetti_paper_figures.

## Acknowledgments

We would like to thank the Vector Institute for Artificial Intelligence for providing the com-puting infrastructure and resources for this work.

## Authors Contributions

G. W. S. conceived the project. G. W. S. and C. M. supervised the project. Z. F. D. developed the SPAGHETTI method, software, and benchmarks. Z. F. D. ran and analyzed benchmarks. Z. F. D. generated experimental results. Z. F. D. ran and analyzed data. Z. F. D. and G. W. S. wrote and edited the manuscript. All authors reviewed the manuscript.

## Supplementary Information

### Supplementary Figures

**Supplementary Figure S1:**
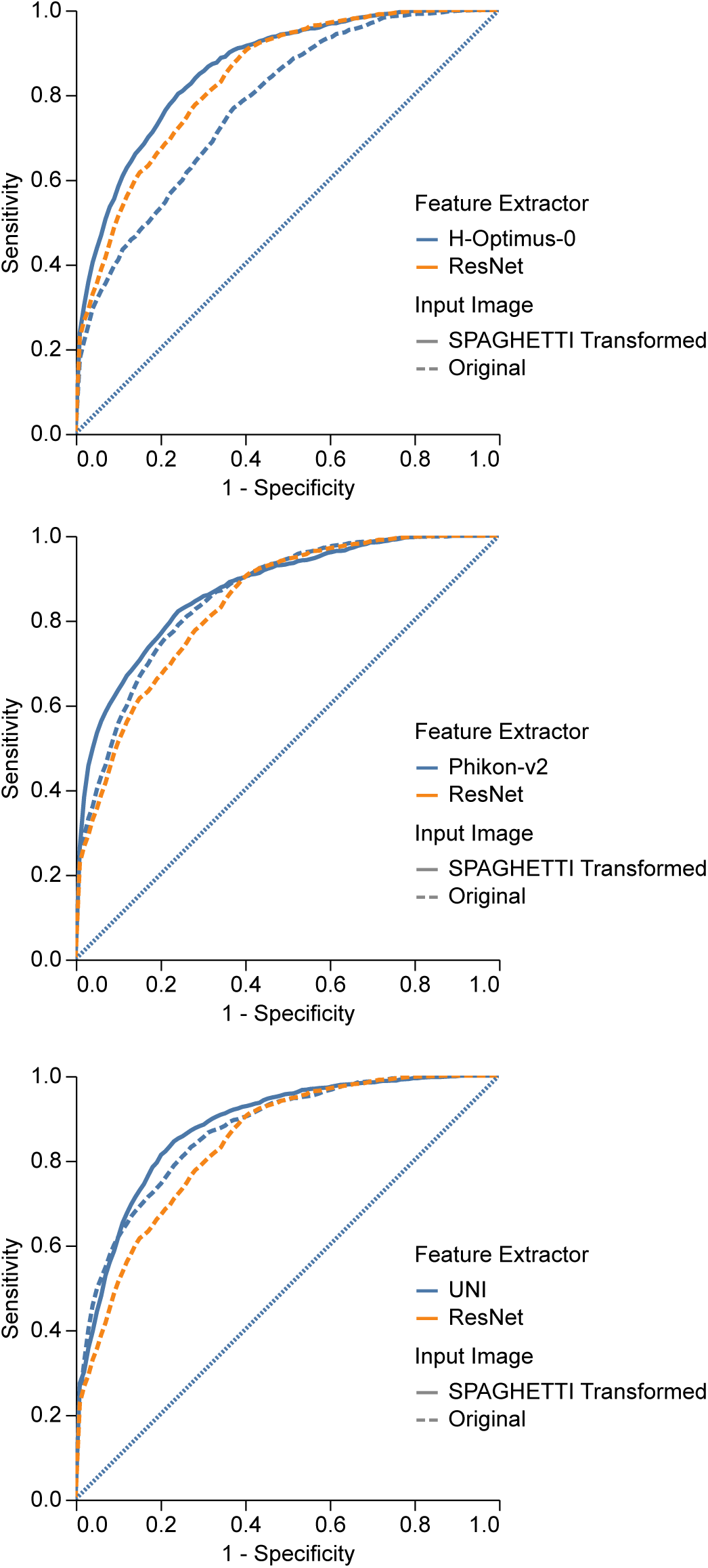
The mean micro ROC curves from 10-fold cross validation for classification of LIVECell cell type, with a random forest classifier on features extracted using H-Optimus-0 (top), Phikon-v2 (middle), and UNI (bottom) from both SPAGHETTI-converted PCM and raw PCM images.

**Supplementary Figure S2:**
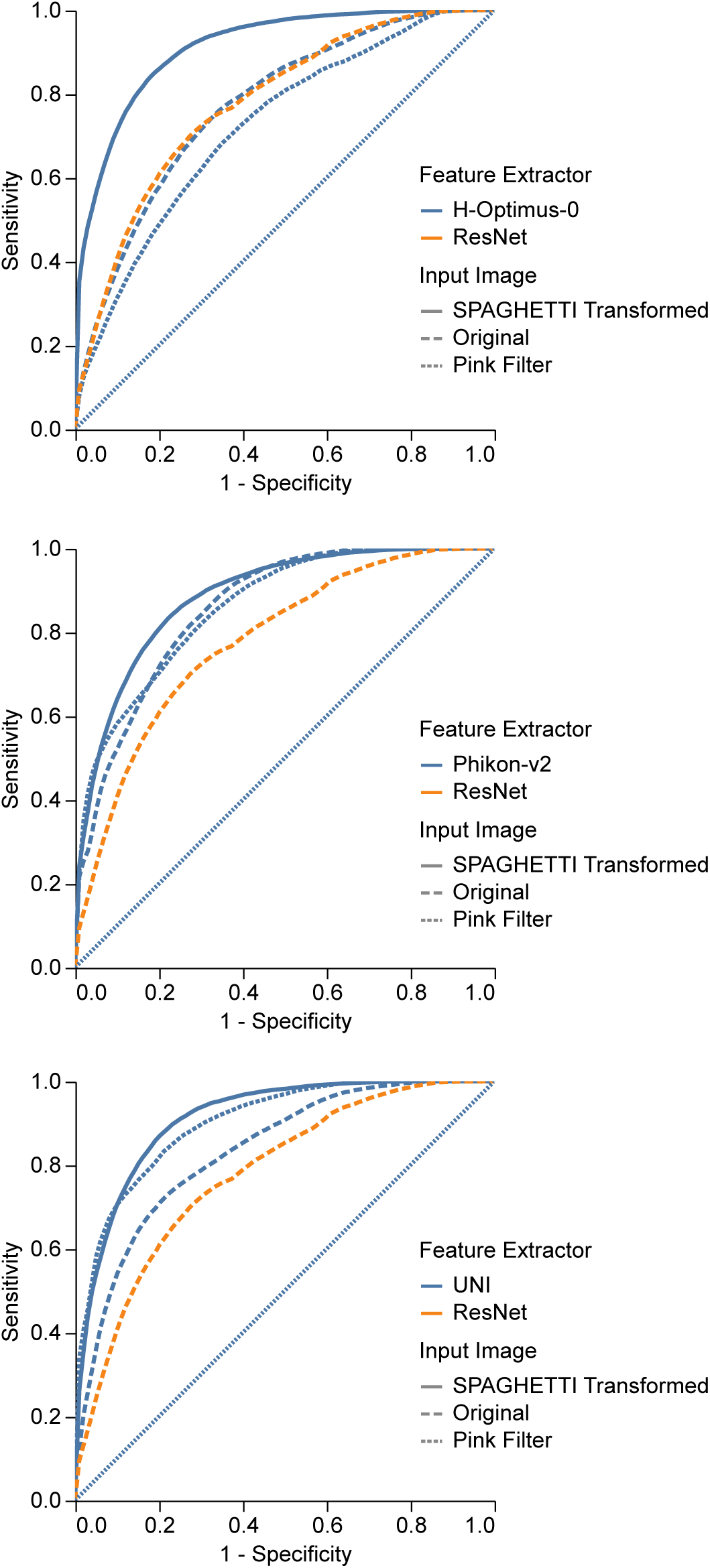
The mean micro ROC curves from 10-fold cross validation for classification of C2C12 media conditions with a random forest classifier on features extracted using H-Optimus-0 (top), Phikon-v2 (middle), and UNI (bottom) from SPAGHETTI-converted PCM, pink-filter transformed PCM, and raw PCM images.

**Supplementary Figure S3:**
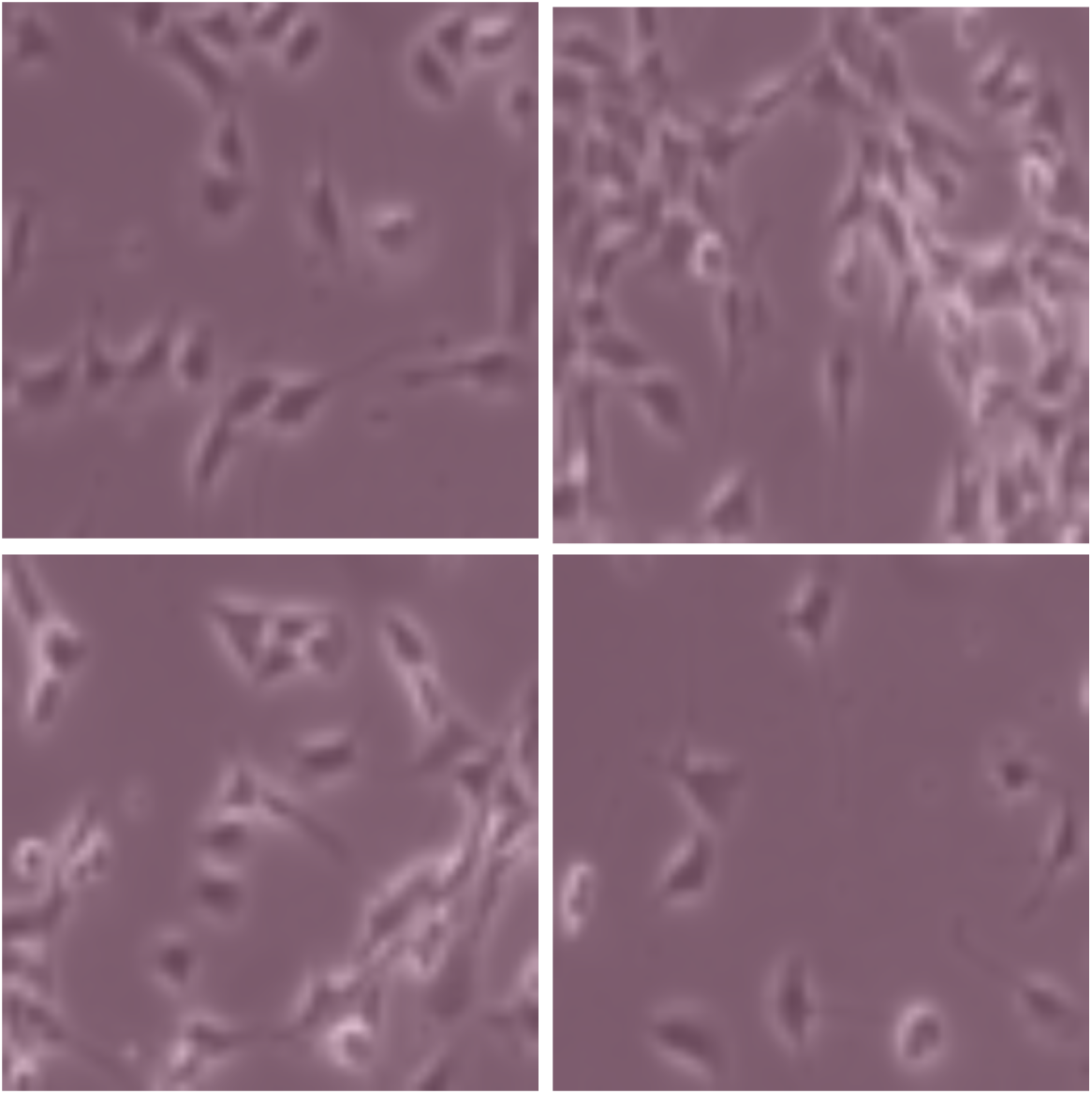
Examples of the pink filter transformed images from the C2C12 dataset.

**Supplementary Figure S4:**
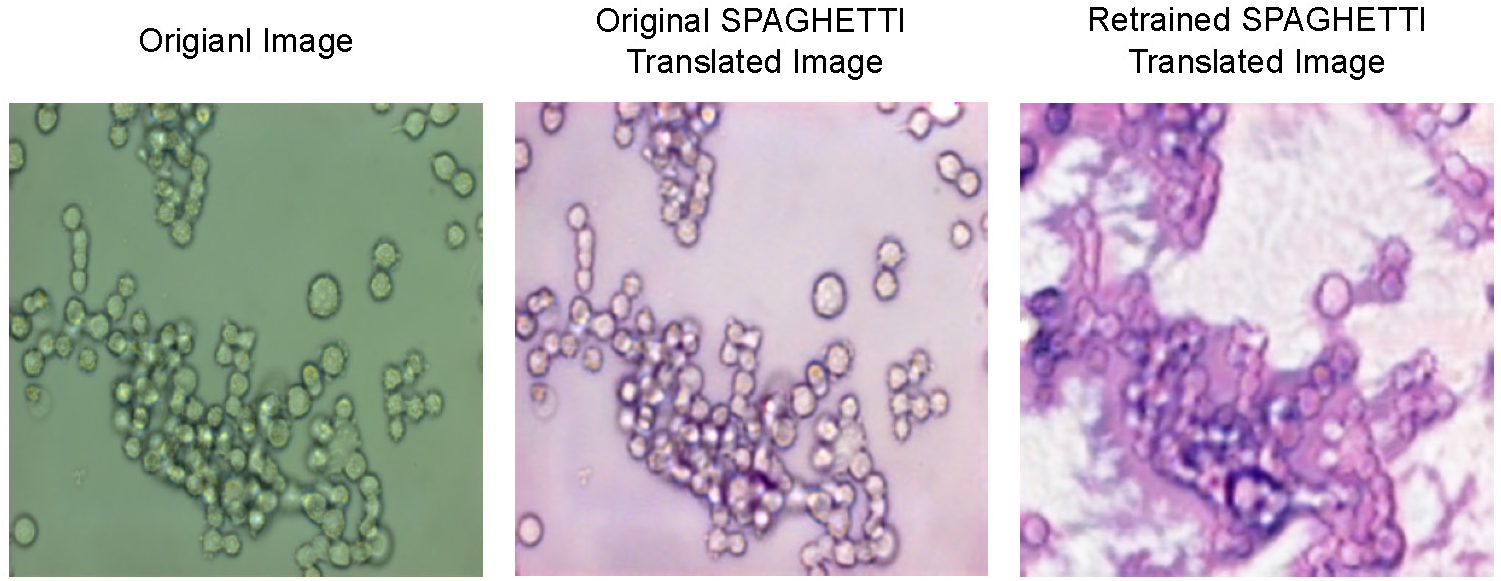
Examples of the original image (left), original SPAGHETTI translated image (middle), and retrained SPAGHETTI translated image (right) from the Caco-2 dataset.

**Supplementary Figure S5:**
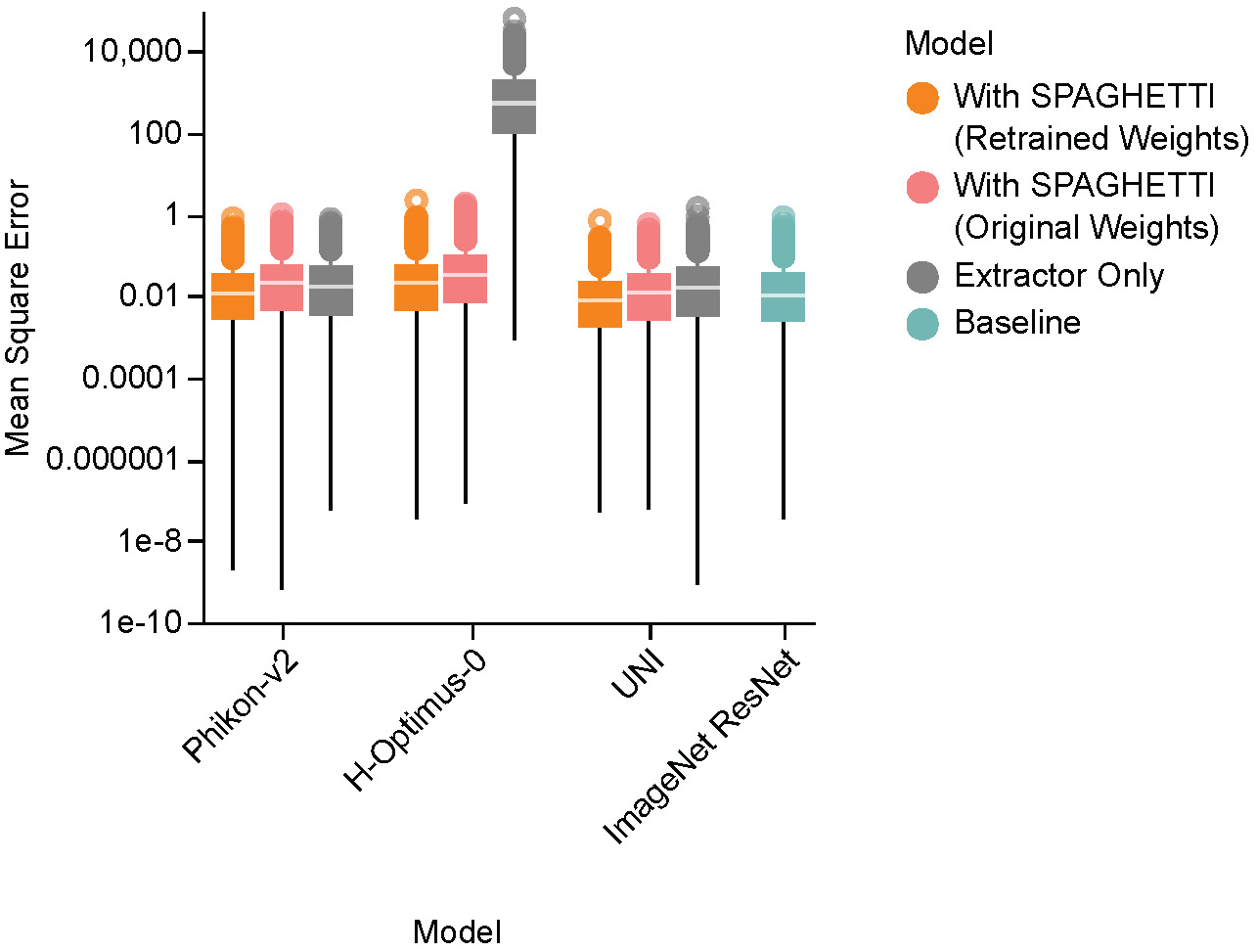
The mean square error between ground-truth and predicted cell viability scores using a linear regression model with retrained SPAGHETTI, original SPAGHETTI, and original PCM images across different feature extractors on the Caco-2 dataset.

**Supplementary Figure S6:**
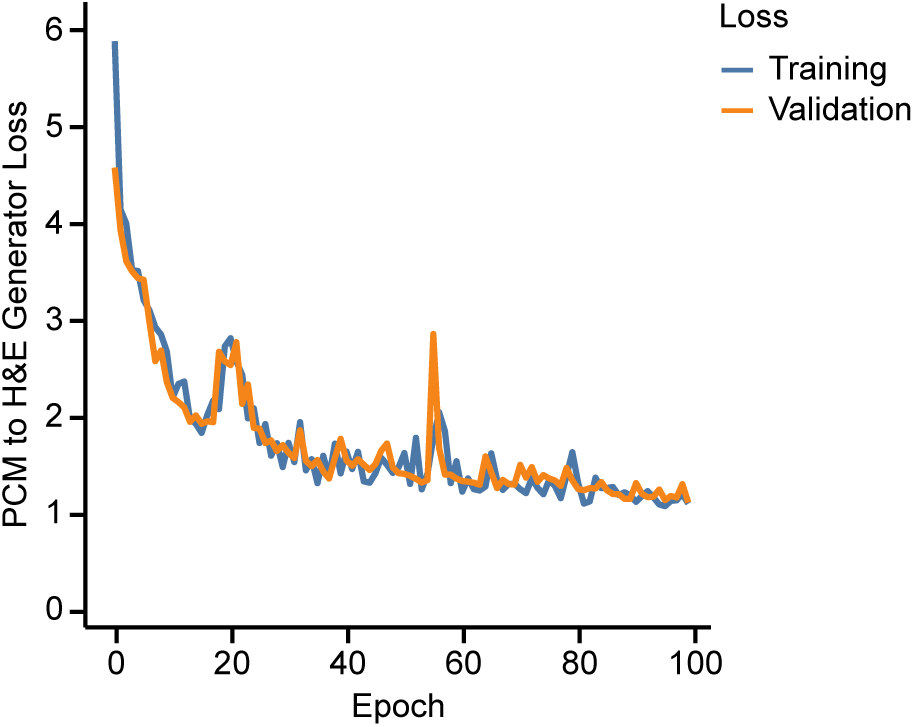
The training and validation loss curves of the SPAGHETTI PCM to H&E generator during the retraining process on the Caco-2 dataset. Retraining is efficient and converges quickly within 100 epochs.

### Supplementary Tables

**Supplementary Table S1:** Training time and dataset size requirements (in number of input images) for training SPAGHETTI and other H&E feature extractor models. Both the original and retrained SPAGHETTI models require much less training time and dataset size compared to other H&E feature extractor models.

| Model | Training GPU Hours (h) | Dataset Size (#) |
| --- | --- | --- |
| SPAGHETTI (Ours) | 128 (NVIDIA T4) | $1.30 \times 10^4$ |
| SPAGHETTI-retrained (Ours) | 72 (NVIDIA T4) | $8.20 \times 10^3$ (300 PCM Images) |
| UNI<sup>11</sup> | 1024 | $1.00 \times 10^8$ |
| Phikon-v2<sup>10</sup> | 4300 (NVIDIA V100) | $4.56 \times 10^8$ |
| H-Optimus-0<sup>9</sup> | N/A | N/A |

